# A geometric atlas of how ESM3 organizes modalities across depth

**DOI:** 10.64898/2026.07.08.737319

**Authors:** Jacob L Steenwyk

**Affiliations:** Department of Molecular and Cell Biology, University of California, Berkeley, Berkeley, 94720, California, USA

**Keywords:** mechanistic interpretability, protein language models, ESM3, multimodal representations, representational geometry, residual stream, protein structure

## Abstract

Protein language models learn general-purpose representations from large collections of protein sequences and structures, and have advanced the prediction of protein structure and function. ESM3 is a multimodal protein language model that ingests a protein through several channels at once, including amino-acid sequence, three-dimensional structure, secondary structure (SS8), solvent accessibility (SASA), and discrete functional annotations, summing their embeddings into a single residual stream. Little is known about whether these modalities occupy separate subspaces and the depth at which they fuse. The present analysis examines ESM3 (esm3-sm-open-v1; 1.4 billion parameters; 48 transformer layers) once per modality in isolation and applies representational-similarity analysis across all 48 layers. The four physical modalities (sequence, structure, SS8, SASA) begin in distinct subspaces, remain maximally separated through roughly the first half of layers, and then fuse into a shared low-dimensional subspace between layers 25 and 35. The fusion is ordered. The structure-derived modalities (structure, SS8, SASA) are mutually aligned from the input, whereas sequence joins last, after layer 28. The functional-annotation modality never fuses; instead, it remains representationally orthogonal to the physical modalities at every layer, and this orthogonality holds whether the annotation is supplied as whole-protein or per-residue, suggesting that it is content-driven rather than a tokenization artifact. The fusion is a learned property, absent in a randomly initialized model of the same architecture, holds at the residue level below the mean-pool, and reorganizes variance, converting between-condition variance into within-condition variance while the stream never approaches isotropy. Fusion depth is independent of protein length but is delayed by structural disorder. The phenomenon is universal across diverse organisms. Across 5,555 proteins from 12 organisms spanning eukaryota, bacteria, and archaea, every superkingdom (and every individual organism) reaches peak modality fusion at the same network depth (layer 35).

## 1 Introduction

Sequence-only protein language models learn structural and functional features from amino acid sequences alone [1]. Multimodal protein language models such as ESM3 [2] reason jointly over sequence and structure. Each input modality has its own embedding table, and the per-residue embeddings are summed before the transformer trunk. This architecture raises a mechanistic question. The modalities may be kept in separate parts of the representation, or the network may blend them. Previous interpretability work on protein language models has examined attention heads and learned features [3–6], but little is known about how the several input modalities of a multimodal model are organized across layer depth.

The present analysis takes a geometry-first descriptive approach to create an atlas of how modalities are organized in representational space. For each protein, ESM3 runs once per modality in isolation, with all other modalities masked, together with a reference of all-modalities, and the residual stream is read out at each layer. The resulting collection of per-(protein, modality, layer) representations forms an atlas of how multimodal information is organized across depth. Standard representational-geometry tools, namely silhouette score, linear CKA, effective rank, and linear probes, characterize the atlas, supported by several controls. This work builds on a previous study, which applied sparse autoencoders to ESM3 and ESM2 and, among other results, reports that the sequence and sequence-plus-structure representations of ESM3 converge between layers 28 and 38 [7]. The present analysis extends that two-condition observation to a full multimodal atlas, characterizing the ordered fusion of the four physical modalities, the persistent orthogonality of the functional channel, the universality of the pattern across diverse organisms from the tree of life, and a battery of controls. Causal steering and sparse-autoencoder dictionaries are developed in companion work [7].

## 2 Results

### 2.1 Physical modalities fuse sharply at mid-depth

The first question is whether the modality conditions occupy separate regions of the representation and, if so, at what depth they merge. To address this question, condition separation was measured at each layer using the silhouette score. Separation increases from 0.32 at the input to a maximum of 0.42 at layer 24, then decreases to a minimum of 0.156 at layer 35, with partial re-separation in the final layers (Figure 1b). The integration index moves inversely. The transition is sharp and localized, with a knee near layer 25 rather than a gradual decline. The accompanying joint-PCA projection tells the same story, showing five distinct clusters in layer 24 that collapse into a single cloud by layer 35, with the exception of functional-annotation (which is discussed in more detail in a later section) (Figure 1a). These metrics converge at the sample size used (Figure S1) and lie far above the label-permutation nulls at each layer (Figure S2).

**Fig. 1:**
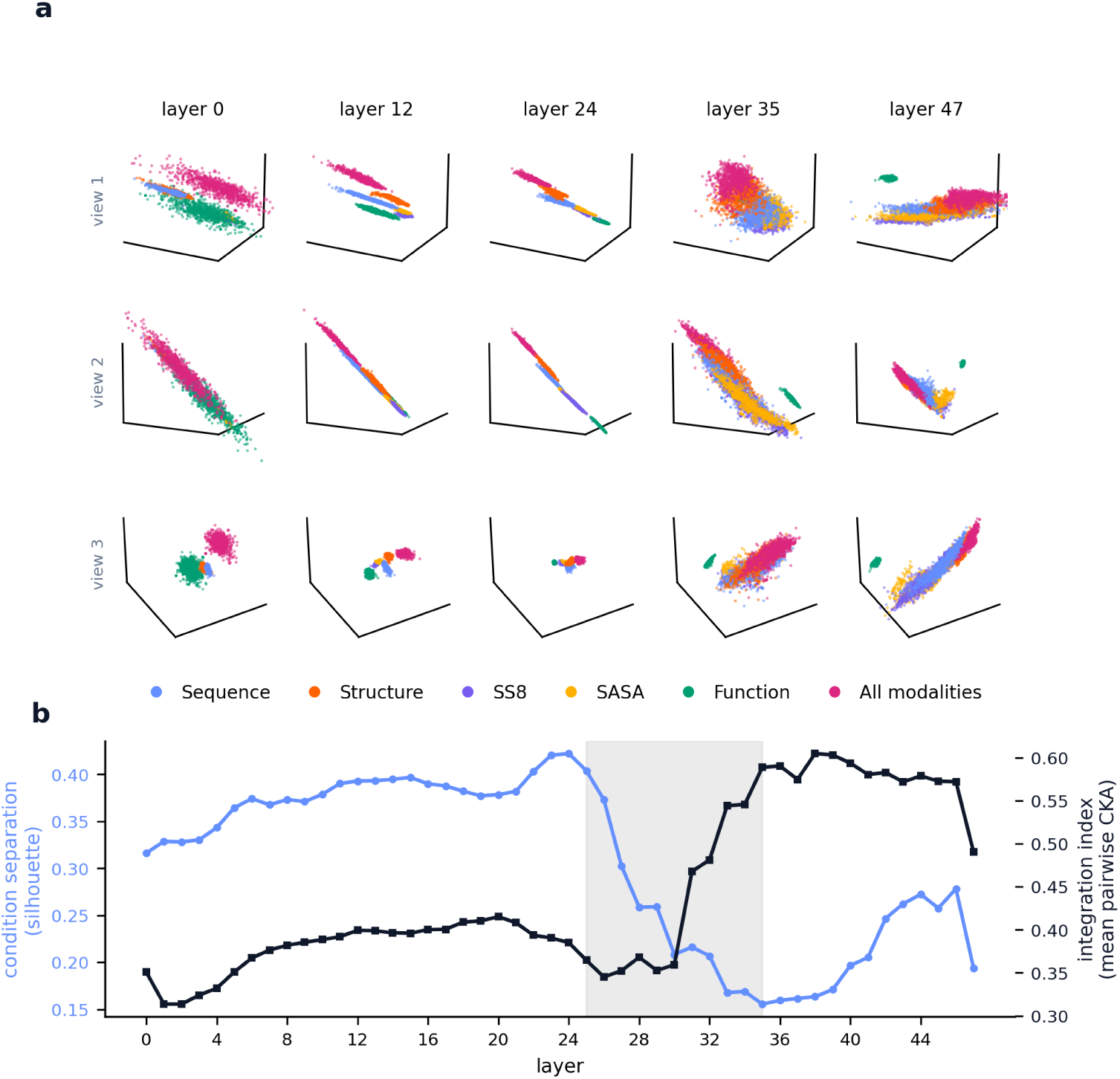
Modality conditions occupy distinct subspaces early and fuse into a shared subspace at mid-depth. (a) Mean-pooled residual-stream representations of 892 human proteins under each single-modality condition, projected with a joint PCA fit across all 48 layers and shown at five depths from three camera orientations (rows), coloured by modality. The five conditions form separated clusters through layer 24 and collapse into one cloud by layer 35, while the functional-annotation condition (green) remains a distinct island. (b) Across all 48 layers, condition separation (silhouette, left axis) rises to a maximum of 0.42 at layer 24, then falls to a minimum of 0.156 at layer 35, while the integration index (mean pairwise CKA, right axis) moves inversely. The shaded band marks the fusion transition (layers 25 to 35).

### 2.2 The fusion is ordered, and sequence joins last

The next question is whether the physical modalities (i.e., those except for functional-annotation) fuse all at once or in a particular order. Per-pair centered kernel alignment (CKA) answers this by tracking, for each pair of conditions, how aligned they become across depth. The structure-derived modalities (structure, SS8, and SASA) are mutually aligned from layer 0, whereas every pair involving sequence stays near a CKA of 0.2 until layer 28 and rises thereafter (Figure 2a; the full condition-by-condition matrix at seven depths is shown in Figure S3). Sequence, the one channel carrying information that cannot be derived from structure, integrates last.

**Fig. 2:**
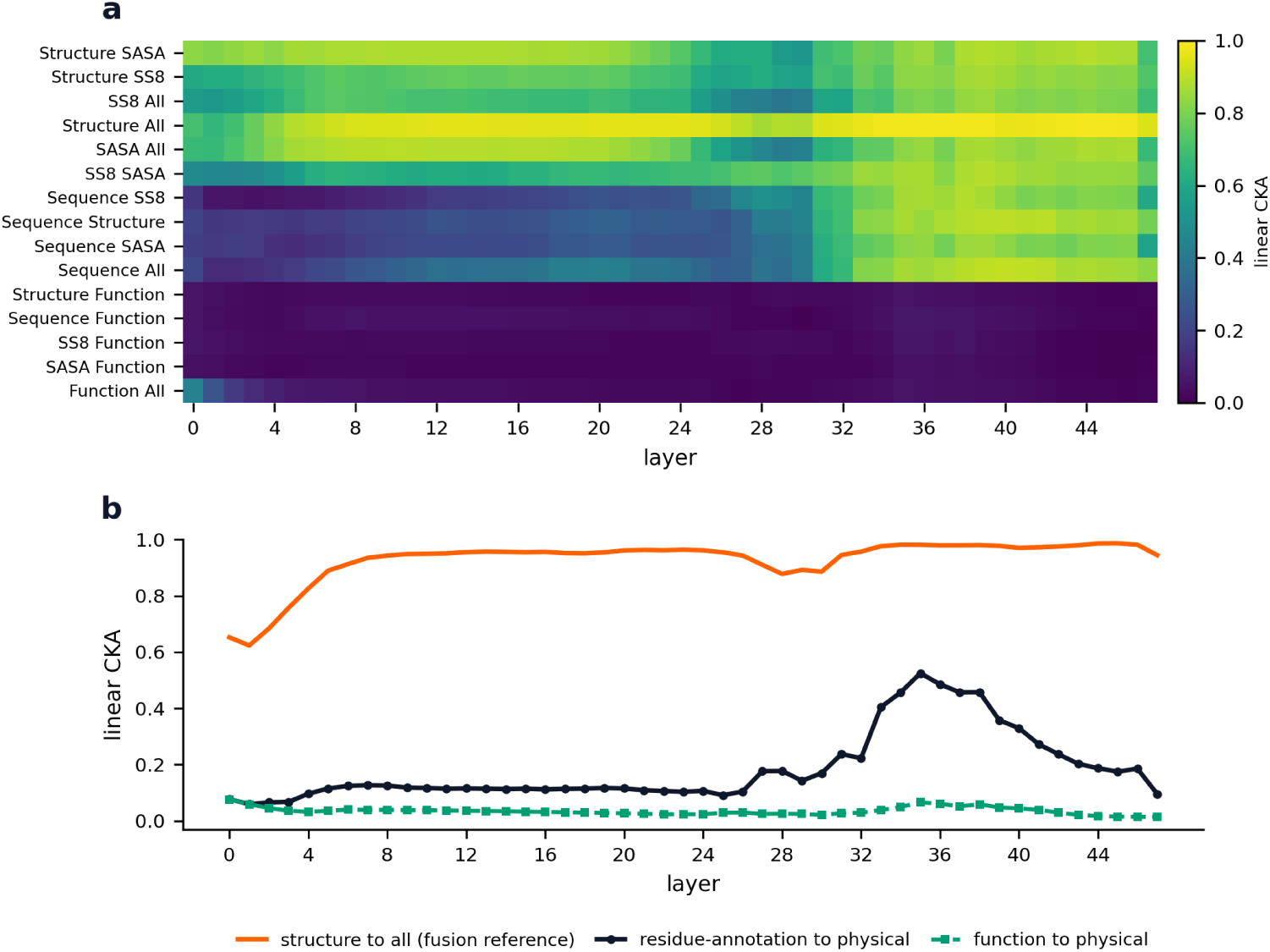
Fusion of the physical modalities is ordered, and the functional channel never joins. (a) Linear CKA between every condition pair at every layer, with rows ordered by fusion onset (the first layer reaching a CKA of 0.5). The structure, SS8, and SASA block is warm from layer 0, the sequence pairs warm only after layer 28, and the four function-versus-physical pairs (bottom rows) stay near a CKA of 0.01 throughout, while the function-to-all pair is warm only at the input, where the all condition still carries the function signal. (b) The whole-protein function track and the per-residue residue-annotation track behave alike. Structure reaches a CKA of about 0.99 with the all-modalities reference (orange), whereas residue-annotation (black) and function (green) both stay below a CKA of 0.55 with the physical modalities.

### 2.3 The functional-annotation modality never fuses

Whether the functional-annotation channel also joins the fusion is a separate question, and it does not. Every pairing of the function condition with a physical modality stays near a CKA of 0.01 to 0.07 across all 48 layers, while structure reaches a CKA of about 0.99 with the all-modalities reference. This orthogonality does not stem from a degenerate representation, because the function cloud keeps genuine cross-protein spread, with an effective rank ranging from 2.8 to 218 across layers. One alternative explanation is granularity, since function is the only whole-protein modality whereas the physical modalities vary residue by residue. To test this explanation, the per-residue annotation modality of ESM3 was added, driven by InterPro residue-site annotations available for 631 of the 892 proteins. The per-residue functional annotation also stays largely orthogonal, with a CKA against the physical modalities near 0.1 and a transient maximum near 0.5, never approaching the CKA of about 0.99 that marks fusion (Figure 2b). Functional information is therefore held in a separate subspace regardless of granularity, a content-driven separation.

### 2.4 The fusion is learned, not architectural

Fusion could, in principle, come simply from summing the modality embeddings, rather than from anything the network learns during training. To tell these apart, the same architecture was run with random, untrained weights in the trunk (the stacked transformer blocks) and the modality embeddings, keeping the trained structure encoder so that the inputs still carry meaning. The random model shows no fusion. Both metrics remain flat across all 48 layers, with condition separation near 0.62 and the integration index near 0.71, and neither changes with depth (Figure 3a). The trained model behaves very differently, with separation that rises and then collapses and an integration index that climbs with depth. The random model’s integration index is high but constant because random projections of the same inputs are always somewhat correlated, unlike fusion. The depth-dependent shift into a shared subspace, therefore, arises from training and is not an automatic consequence of simply adding the modality embeddings.

**Fig. 3:**
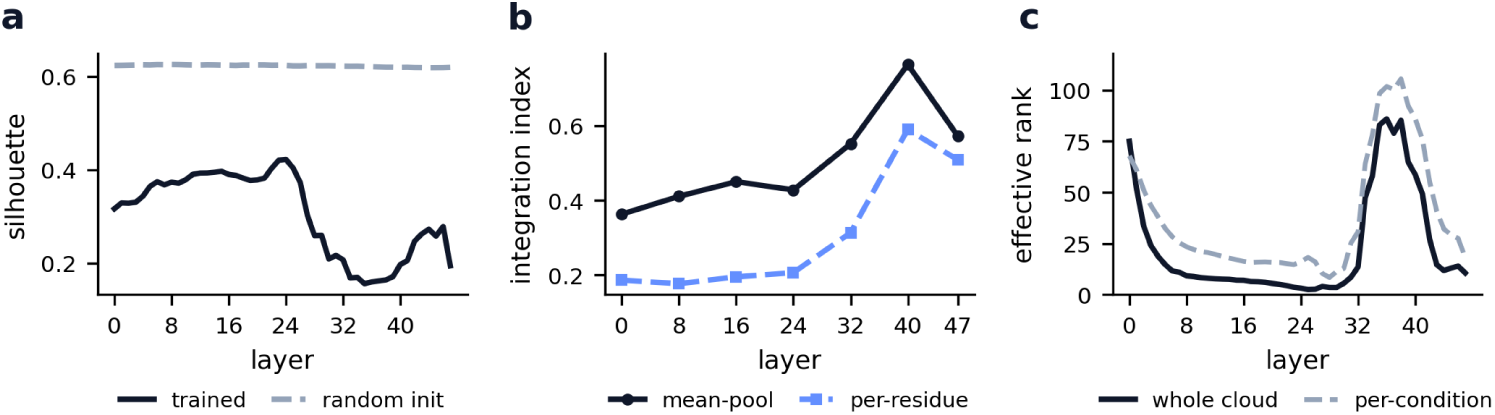
The fusion is learned, holds below the mean-pool, and reorganises variance without approaching isotropy. (a) A model with the same architecture but randomly initialised weights shows no fusion, with condition separation near 0.62 at every layer against the trained model’s peak-then-collapse. (b) On residue-level representations of a 100-protein subset, the integration index rises with depth (0.18 to 0.59), mirroring the pooled curve. (c) Effective rank of the whole cloud collapses to about 3 at the layer-24 separation peak, expands to about 85 of 1536 in the fusion zone, then contracts, and the whole-cloud and per-condition ranks converge once fused.

### 2.5 Fusion holds below the mean-pool

The main analysis averages each protein’s residual stream over its residues, which raises the hypothesis that this averaging creates the appearance of fusion. To evaluate this, the integration index was measured again in individual residue representations, without averaging, for a 100-protein subset across seven layers of the network. The residue-level integration index still increases with depth, from 0.18 at the input to 0.59 in the fusion zone, following the same upward path as the pooled curve (Figure 3b). The convergence is therefore already present in the individual residue representations and is not produced by the averaging step. Single residues, and not only whole-protein averages, move into a shared subspace at mid-depth, so fusion is a real property of the representation rather than a side effect of how it is summarized.

### 2.6 Fusion reorganizes variance without approaching isotropy

When the conditions fuse, it is not clear what happens to the representation’s shape. For example, fusion could spread the stream out into a diffuse, high-dimensional cloud or collapse it toward a single point. To examine this, two quantities were tracked as a function of depth. The first is the effective rank of the stream, a measure of how many dimensions carry real variance. The second is the accuracy of a probe that tries to read the modality identity back out of the stream. The effective rank of the whole cloud collapses to about 3 at the layer-24 separation peak, then expands to about 85 of a possible 1536 in the fusion zone before contracting, and the whole-cloud and per-condition ranks converge once the conditions are fused (Figure 3c). Therefore, fusion shifts the variance from between conditions to within each condition, without ever allowing the stream to spread toward isotropy (i.e., the same spread in every direction). Modality identity is not lost either. A logistic probe reaches an accuracy of 1.0 in the full 1536-dimensional stream at every layer, so a thin additive identity signal always remains, whereas the same probe in the top-3 PCA geometry falls to about 0.68 in the fusion zone, driven mainly by SASA, even as the function condition stays decodable throughout (Figure S4). These findings suggest that fusion is a controlled reorganization of variance rather than a collapse or a spreading out of the representation.

### 2.7 Fusion depth tracks, albeit weakly so, with secondary-structure content

Across the 892 proteins, the depth of fusion-onset is independent of protein length (Spearman r of −0.04, *p* = 0.26) but is delayed by structural disorder, with a coil-fraction correlation of +0.22 (*p* = 3 × 10^−11^) and a helix-fraction correlation of −0.20 (*p* = 3 × 10^−9^), so that well-folded helical proteins fuse earlier (Figure 4a,b). However, no correlation was observed between strand-fraction and fusion-onset layer (*r* = 0.04, *p* = 0.28). Fusion-onset has no relationship with AlphaFold confidence (mean correlation of pLDDT of +0.003, p of 0.93). The disorder effect may partly stem from secondary-structure content rather than the model’s structural uncertainty. The same ordering is visible across the ten categories of curated proteins, from receptors and small proteins that fuse earliest to DNA-binding proteins that fuse latest (Figure S5).

**Fig. 4:**
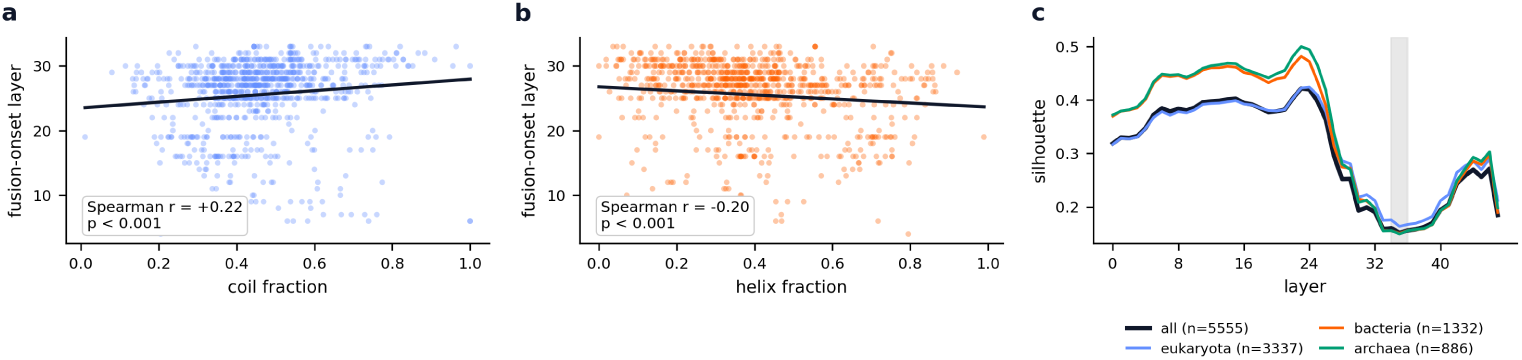
Fusion depth tracks secondary-structure content and is universal across the tree of life. (a) Per-protein fusion-onset layer averaged within each of ten protein categories (bars, mean and standard error). Onset is delayed by structural disorder (Spearman r of +0.22 against coil fraction) but is unrelated to protein length (*r* = −0.04, n.s.) or to AlphaFold confidence (mean pLDDT *r* = +0.003, n.s.). (b) Condition separation across depth for 5,555 proteins from 12 organisms, stratified by superkingdom. The curves are nearly superimposable and reach minimum separation at layer 35 (shaded); every one of the 12 organisms reaches its minimum at layer 35.

### 2.8 Fusion is universal across diverse organisms in the tree of life

Across 5,555 proteins from 12 organisms that span all three superkingdoms (Figure S6), the pooled fusion curve is nearly identical to the human-only result, with a silhouette maximum of 0.42 at layer 23 and a minimum of 0.152 at layer 35 versus 0.42 at layer 24 and 0.156 at layer 35. Stratified by superkingdom, the eukaryota, bacteria, and archaea curves show the same peak-then-collapse and reach minimum separation in layer 35 (Figure 4c), with maxima of 0.42, 0.48, and 0.50 and minima of 0.164, 0.151, and 0.149. Each of the 12 organisms reaches its minimum in layer 35, with a standard deviation of 0 (Figure S7). Superkingdoms are intermixed in the representation rather than separated, so the organization is by modality and not by source organism (Figure S8). Multimodal fusion is therefore a universal, depth-locked property of ESM3 and not an artifact of the curated human set.

## 3 Discussion

ESM3 organizes its inputs into two representational regimes. The four physical modalities collapse into a shared, low-dimensional subspace at mid-depth, whereas the functional-annotation modality occupies a subspace that stays orthogonal throughout the network. The mid-network timing of physical fusion, the late entry of sequence, and the dependence of fusion depth on secondary-structure content together suggest that the network first builds modality-specific features and then commits to a unified physical representation once secondary structure is resolved, while holding discrete functional labels on a separate axis. The depth-locked universality across diverse organisms from the tree of life indicates that this organization is intrinsic to the trained model rather than a property of any particular proteome.

Geometric orthogonality does not mean that functional content is absent from the fused representation. A linear probe recovers the enzyme class from the structure-derived representation with a balanced accuracy that rises from 0.56 to about 0.79 by layer 39, and it recovers the fold class from the physical conditions at about 0.84 throughout (Figure S9). The explicit function channel behaves in the opposite way, decoding the fold class only near 0.40 and the enzyme class near 0.55, the latter declining in depth. The network, therefore, extracts functional content into the physical subspace while keeping the discrete function channel on a separate, informationally thin axis, so that the orthogonality is a representational choice rather than an absence of functional information. A causal intervention supports this notion. Withholding the function input from the full model displaces the fused representation about six times less than withholding structure, sequence, or SASA, with a median relative displacement of 0.02 against 0.07 to 0.12 at the fusion layer (Figure S10), so the fused physical representation is largely insensitive to the function input.

One mechanism that could keep functional annotation orthogonal is redundancy. Because the enzyme class is decodable from the structure-derived representation, the network may already recover function from the physical modalities and face little pressure to integrate the explicit functional channel. This account predicts that the function channel should align more closely with the physical subspace for proteins whose functions cannot be recovered from the structure. The prediction fails (Figure S11). The alignment Per-protein between the function vector and the physical sub-space is near zero for every protein, with a mean of 0.0005 and a standard deviation of 0.15, and it does not track redundancy. Proteins for which the structure representation misclassifies enzyme class align no more strongly than proteins it classifies correctly, with means of 0.014 and 0.006 and a Mann-Whitney *p* of 0.35, and alignment is flat against both the number of InterPro domains, with a Spearman correlation of −0.02, and the number of gene-ontology terms, with a Spearman correlation of 0.09. Functional annotation, therefore, stays orthogonal whether or not it is redundant with the physical modalities, which points away from redundancy and suggests a categorical organization in which discrete functional labels are held on a separate axis irrespective of their recoverability from structure.

Several limitations bound these conclusions. Only one model was studied, because esm3-sm-open-v1 is the sole openly available multimodal ESM3, larger ESM3 models are reachable only through a gated interface, and the ESM2 family [8] is sequence-only and cannot support the experiment. Whether the fractional fusion depth scales with model capacity therefore remains open. The main structures are AlphaFold predictions, but the fusion signature is not an artifact of predicted coordinates, because the disorder effect is uncorrelated with prediction confidence and the silhouette dip at layer 35, the integration-index rise, and the orthogonality of the function channel all reproduce on 177 proteins with experimental coordinates (Figure S12). Finally, the present work characterizes the geometry of the residual stream and does not intervene on it, so the analysis is descriptive rather than causal.

## Methods

### Model and modalities

The model used throughout is esm3-sm-open-v1 [2] (1.4 billion parameters, 48 transformer blocks, model dimension 1536). Six conditions are studied, comprising five single-modality inputs (sequence, structure, SS8, SASA, and function) and an all-modalities condition in which every modality is supplied jointly. Structure is provided as vector-quantized structure tokens derived from coordinates, SS8 is computed by DSSP [9] and SASA by the Shrake-Rupley algorithm [10] from the structure, and function is provided as whole-protein InterPro annotations [11] filtered to the 29,026-entry vocabulary of the ESM3 function tokenizer.

### Datasets

Three datasets were used, namely a 199-protein human pilot for development, an 892-protein human set for the main results, and a set of 5,984 proteins curated from 12 organisms spanning eukaryota, bacteria, and archaea, of which the 5,555 with complete structure and annotation were analyzed, for the universality test. Structures are AlphaFold-DB models of a single release [12, 13], SS8 and SASA are computed with DSSP (mkdssp 4.6.1, via ESM’s ProteinChain) [9], and InterPro and Gene Ontology annotations are taken from UniProt [11, 14]. A fourth dataset of 177 proteins backed by experimental (X-ray and cryo-EM) structures from the RCSB replicates the main result with non-predicted coordinates.

### Harvesting

For each (protein, condition) pair, ESM3 runs one forward pass and the residual stream is cached at all 48 layers, mean-pooled over residues with the special tokens excluded. A 100-protein subset additionally stores per-residue activations at seven layers for the pooling control.

### Metrics

Each layer is summarized by the silhouette score [15] of the conditions (Euclidean, in the full 1536-dimensional space), by the linear CKA of Kornblith et al. [16] between every protein-aligned condition pair, by an integration index defined as the mean pairwise CKA, by the effective rank [17] computed as the exponential of the entropy of the covariance eigenspectrum for the whole cloud and for each condition, and by a protein-grouped five-fold logistic probe of modality identity evaluated both in the full space and in the top-3 PCA geometry. Visualizations use PCA, fit per layer and in a shared basis across layers.

### Controls

The controls comprise a randomly initialized model whose trunk and modality embeddings are reset while the structure encoder is preserved, a per-residue replication, a subsampling-convergence analysis, label-permutation nulls, and protein-bootstrap confidence intervals.

## Data availability

The input data are public. Structure models are from AlphaFold-DB [13] and experimental structures from the RCSB, and sequence, InterPro, and Gene Ontology annotations are from UniProt [11, 14]. Accessibility lists for every data set, including the exact RCSB entries used for the experimental replication, are available in the GitHub repository (https://github.com/JLSteenwyk/esm3-modality-atlas). The mean bundled residual-stream activations are deposited on figshare (DOI: 10.6084/m9.figshare.32928713).

## Code availability

All analysis code is available at https://github.com/JLSteenwyk/esm3-modality-atlas. The pipeline is configuration-driven and provides, as separate scripts, dataset curation; structure retrieval from AlphaFold-DB and the RCSB; secondary-structure and solvent-accessibility annotation; activation harvesting; leave-one-out modality ablation; aggregation; the metric and control analyzes; and figure assembly. Random seeds are fixed throughout.

Representations were extracted from esm3-sm-open-v1 [2], accessed through the EvolutionaryScale esm package (version 3.2.1) under its open license. The main dependencies are Python 3.10, PyTorch 2.6 [18], scikit-learn 1.7 [19], SciPy 1.15 [20], and NumPy 1.26 [21]. The secondary structure is computed with DSSP (mkdssp 4.6.1) [9] and solvent accessibility with the Shrake-Rupley implementation [10] in ESM’s ProteinChain. Figures use the colorblind-safe pypubfigs palette, a Python port of ggpubfigs [22].

## Acknowledgements

The author thanks the developers of ESM3 for making the model publicly available.

## Supplementary Figures

**Figure S1.**
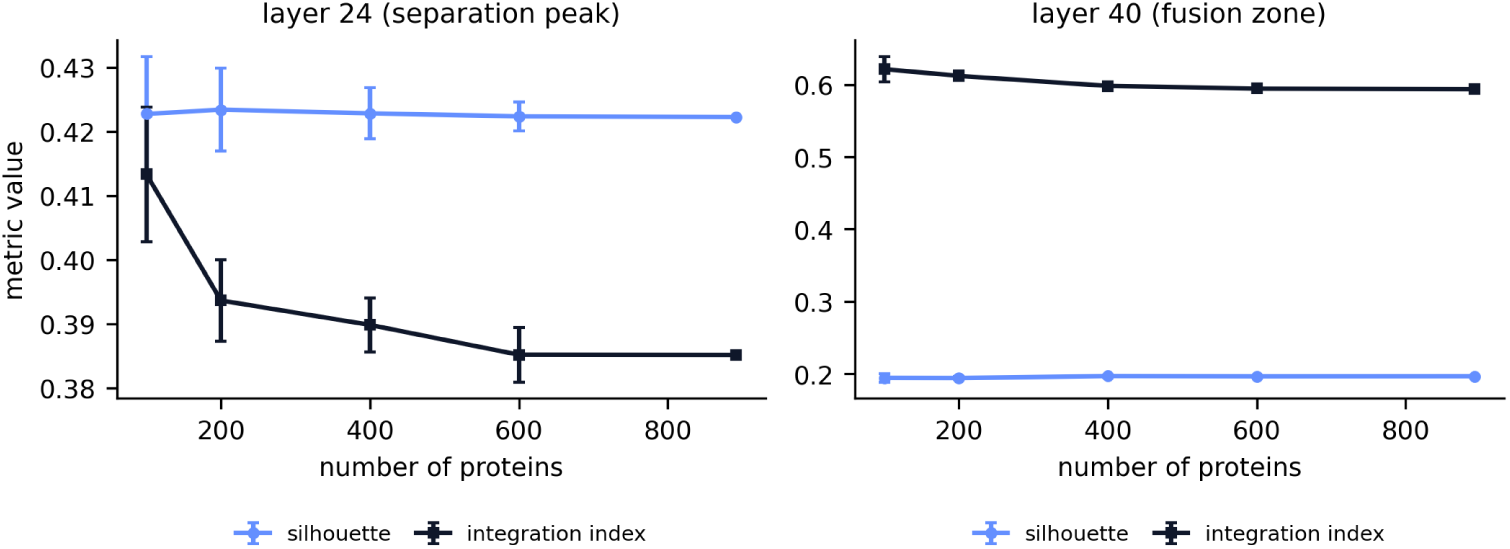
The metrics are converged in the sample size used, reflected in the silhouette and integration index calculated on random subsamples of the 892-protein set (mean and standard deviation over 12 draws per size) at layer 24 (separation peak) and layer 40 (fusion zone). The silhouette is stable from 100 proteins, and the integration index settles by 400 to 600 proteins, so 892 proteins lie within the converged regime for every reported quantity.

**Figure S2.**
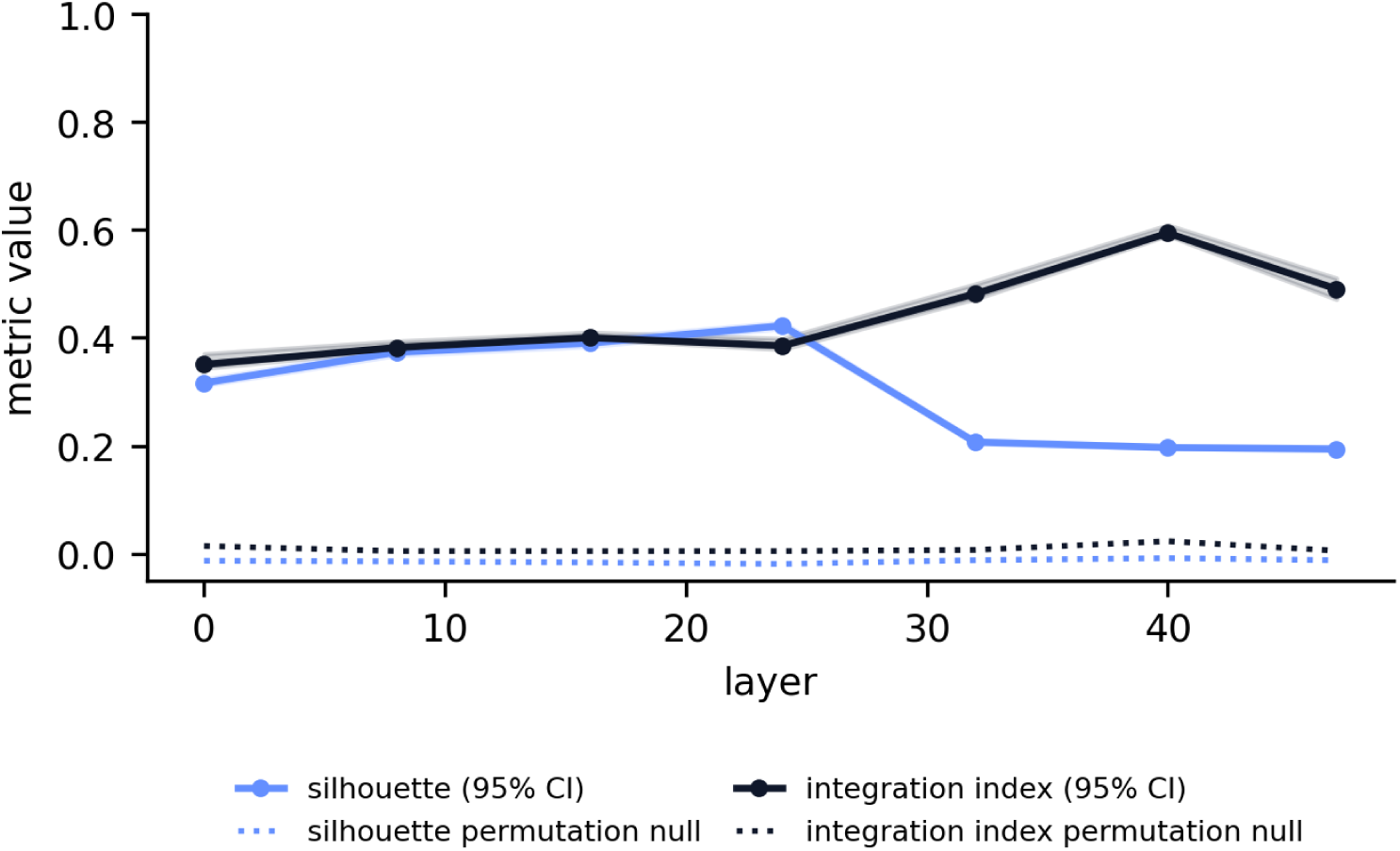
Every fusion metric is significant against a permutation null; For the canonical layers, the observed silhouette and integration index (points, with 95percent confidence bands of the protein boot) sit far above their label-permutation null means (dotted), with a permutation p of 0.005 at every layer tested.

**Figure S3.**
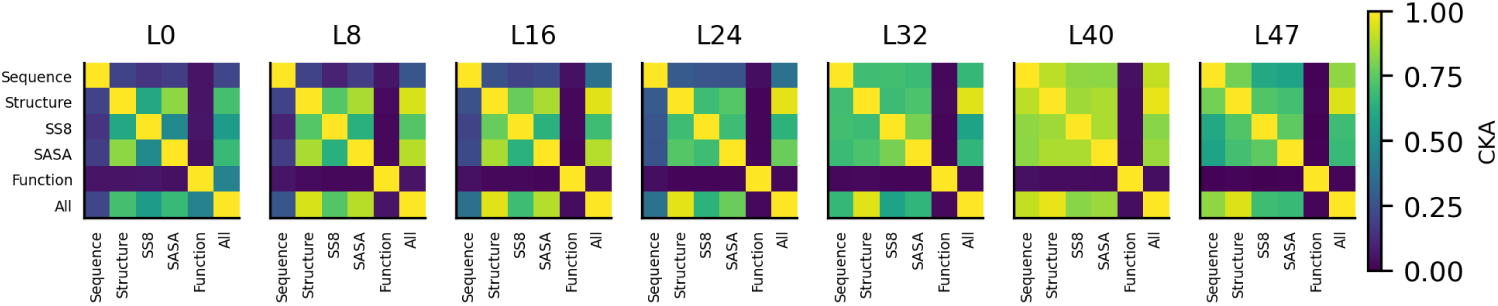
The condition-by-condition CKA matrix warms with depth except along the function axis. Linear CKA between all six conditions at seven depths. The structure, SS8, and SASA block is warm from layer 0, the sequence entries warm after layer 24, and the function row and column stay near CKA 0 at every depth.

**Figure S4.**
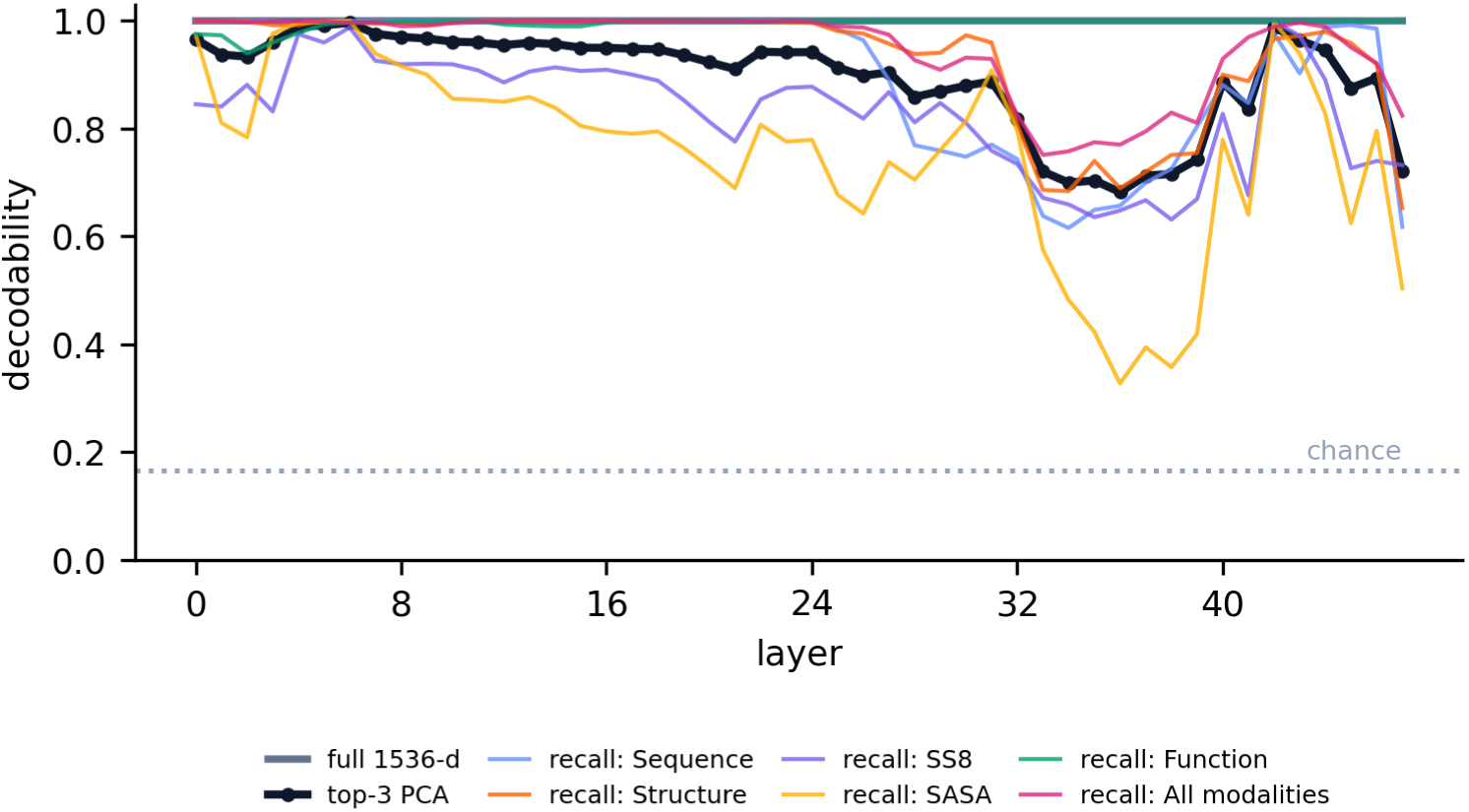
Modality identity remains decodable, and the functional channel remains decodable longest. A protein-grouped logistic probe of modality identity. In the full 1536-dimensional stream, decodability is 1.0 at every layer (an additive modality tag persists). In the top-3 PCA geometry, overall accuracy falls to about 0.68 in the fusion zone, driven by SASA (per-condition recall near 0.33), whereas the function condition stays near 1.0 throughout. The chance is 1 of 6.

**Figure S5.**
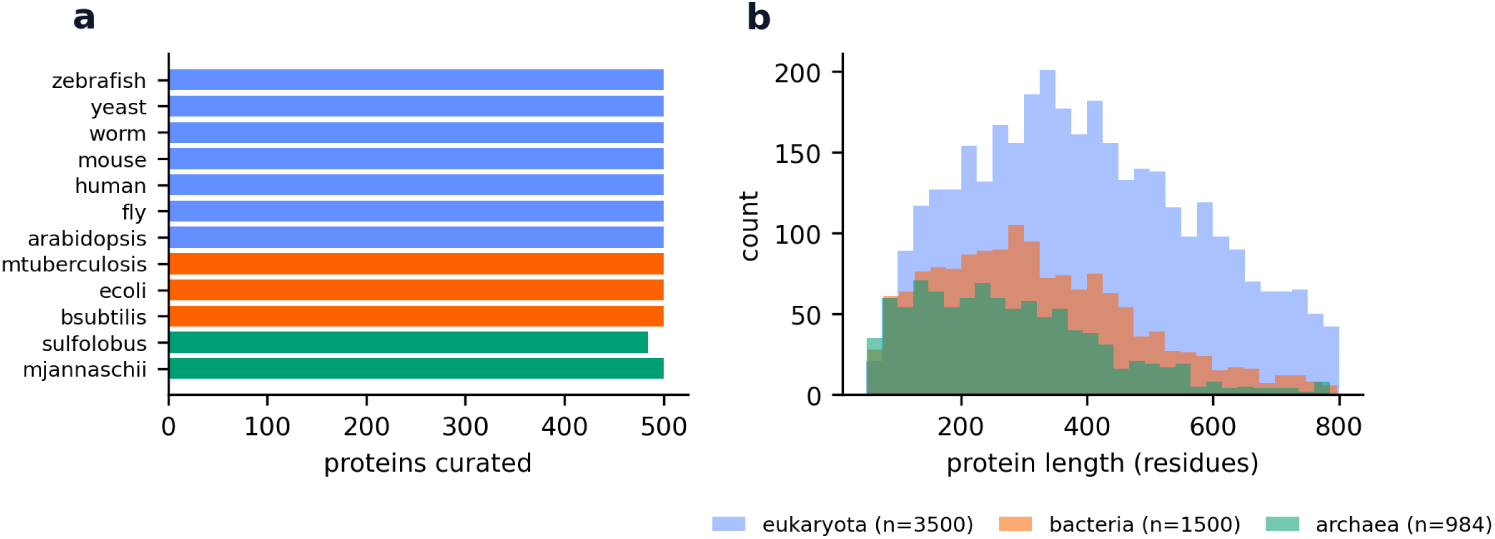
Composition of the diverse set of 12-organisms. (a) Proteins curated per organism (about 500 each), colored by superkingdom. (b) Protein-length distributions by superkingdom over the 50 to 800 residue range.

**Figure S6.**
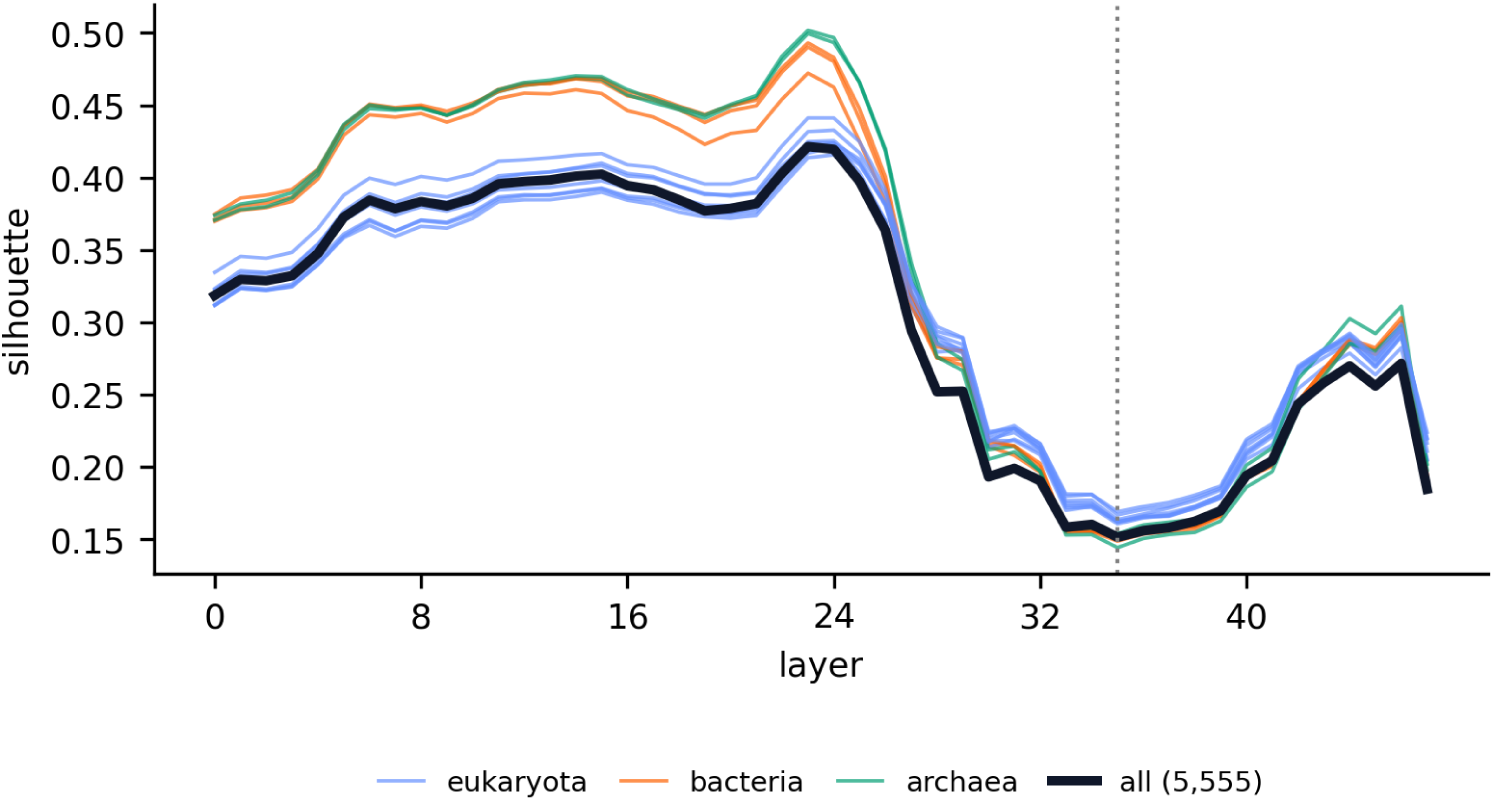
Every individual organism reaches peak fusion at the same depth. Condition separation across depth for each of the 12 organisms (thin lines, colored by superkingdom) and for the pooled set (thick line). All 12 organism curves and the pooled curve reach minimum separation at layer 35 (dotted line).

**Figure S7.**
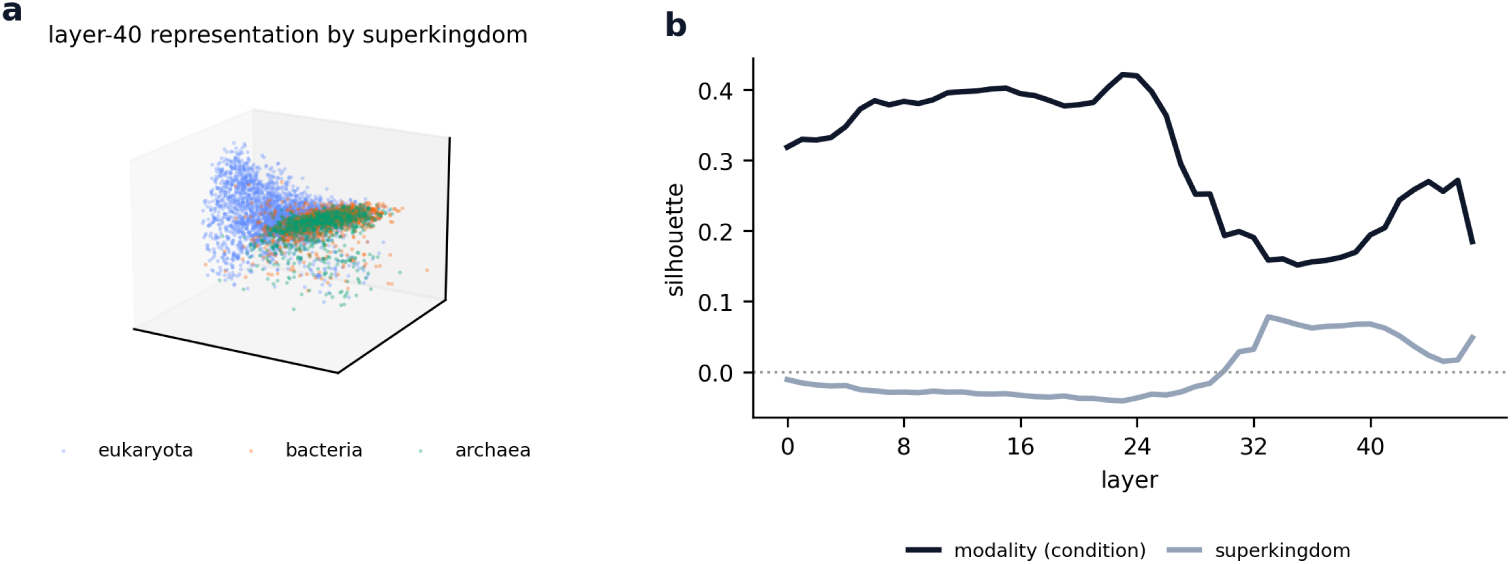
The representation is organized by modality, not by source organism. (a) layer-40 all-modality representation of 5,555 proteins, projected to three dimensions and colored by superkingdom. Eukaryota, bacteria, and archaea are intermixed rather than separated. (b) Across depth, condition separation by modality (silhouette, reaching 0.42 at layer 23) far exceeds separation by superkingdom, which stays near 0 at every layer (peaking at only 0.08 at layer 33). Therefore, ESM3 represents proteins by their biochemistry rather than taxonomy.

**Figure S8.**
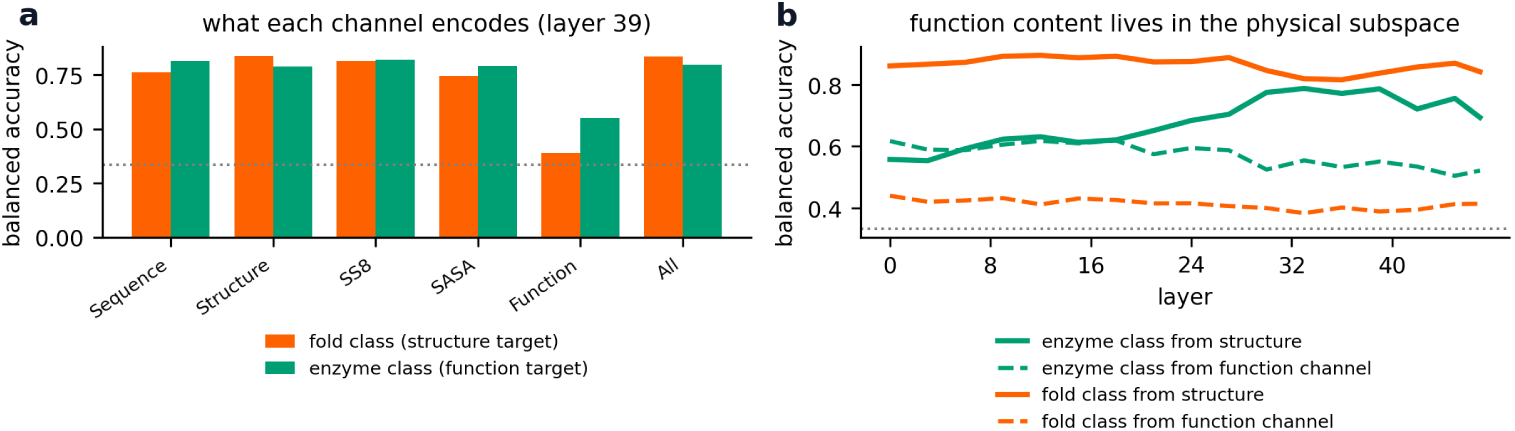
Function content lives in the physical subspace even though the function channel is geometrically orthogonal. A probe (standardization, 50-component PCA, logistic regression, five-fold balanced accuracy) decodes a structural target (fold class from SS8, three classes) and a functional target (EC enzyme class, three classes) from each modality condition. (a) At the post-fusion layer 39, the physical conditions decode both targets well, whereas the function channel decodes both weakly, so there is no division of representational labor. (b) Across depth, enzyme class becomes more decodable from the structure-derived representation, reaching about 0.79, while the function channel decodes fold class near 0.40 and enzyme class near 0.55. Functional information is therefore present in the physical subspace, and the geometric orthogonality of the function channel reflects how the network organizes the representation rather than an absence of functional content. The dotted line marks the chance rate of 0.33.

**Figure S9.**
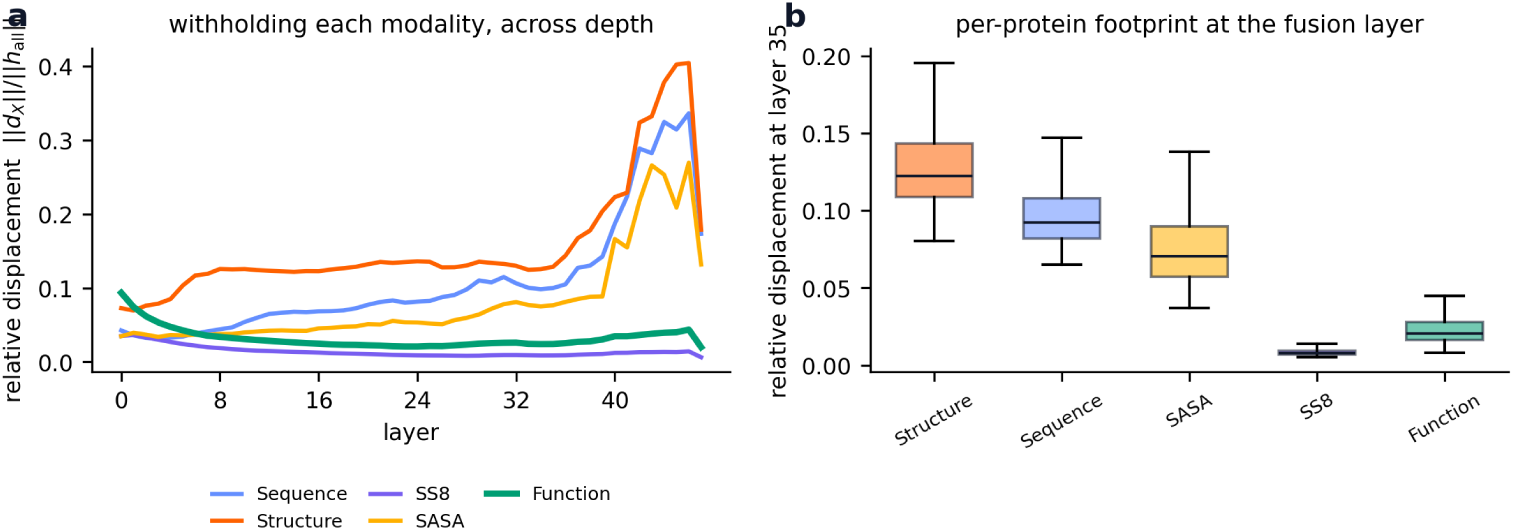
Withholding function barely perturbs the fused representation. Each modality’s contribution is the displacement between the all-modalities representation and the representation with that modality withheld, measured for all 892 proteins as a fraction of the all-modalities vector norm. (a) Relative displacement across depth. Withholding function (green) moves the representation far less than withholding structure, sequence, or SASA, and the gap widens with depth. (b) Per-protein relative displacement at the fusion layer 35. The function footprint, with a median of 0.021, is about six times smaller than the structure footprint, with a median of 0.12. The SS8 footprint is smaller still, but because SS8 is redundant with structure rather than orthogonal, so a small footprint marks low marginal impact, which the geometric and decoding results attribute to orthogonality in the case of function. This is causal evidence that the fused physical representation is largely insensitive to the function input.

**Figure S10.**
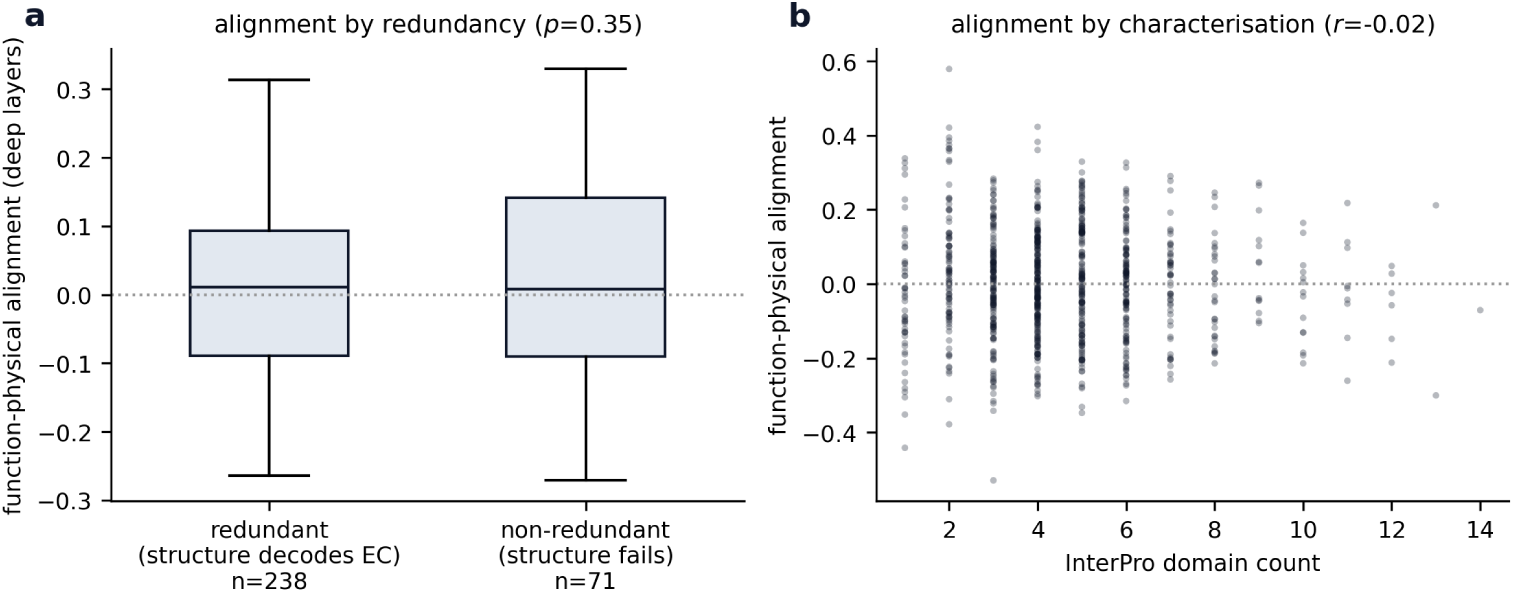
Function orthogonality does not reflect redundancy. Per-protein alignment is the cosine between a protein’s function vector and its mean physical vector, averaged over layers 32, 40, and 47, after centering each condition. If function stayed orthogonal only because it is recoverable from the physical modalities, alignment should be higher for proteins whose function is not recoverable. (a) Proteins split by whether the structure-derived representation correctly decodes enzyme class. The non-redundant group, for which structure fails, aligns no more than the redundant group (medians near 0, Mann-Whitney p = 0.35). (b) Alignment against InterPro domain count shows no trend (Spearman r = −0.02). Alignment is near zero for every protein regardless of redundancy, which favors a categorical account of the functional channel over a redundancy account.

**Figure S11.**
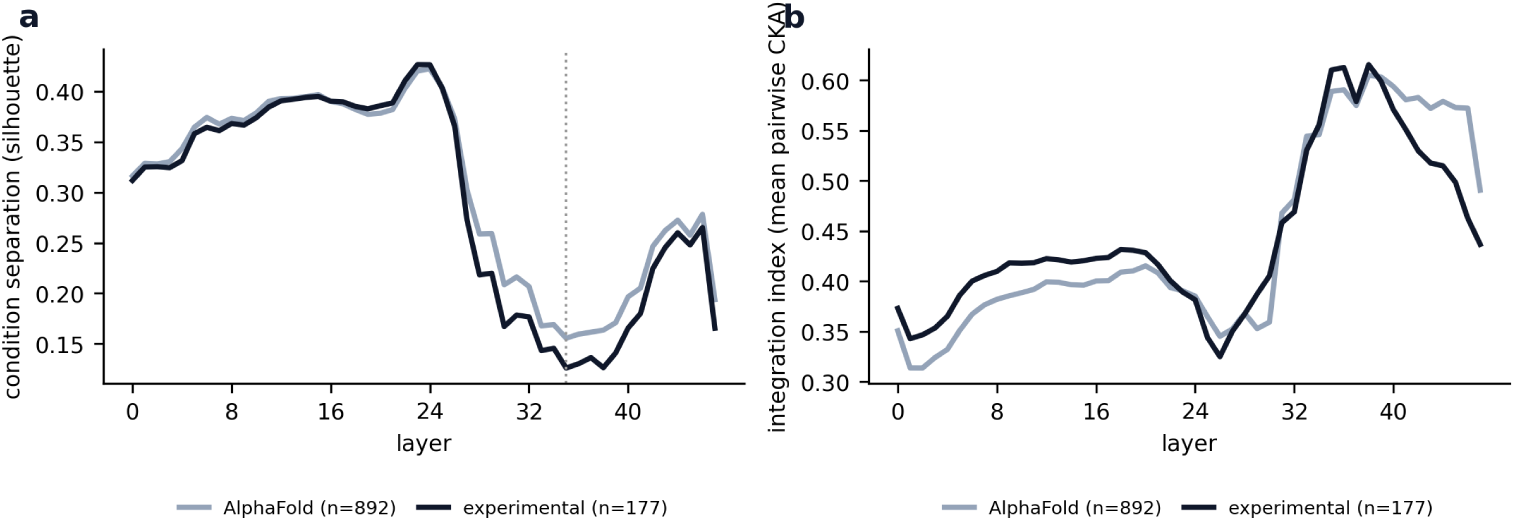
The fusion signature replicates in experimental structures. The main analysis was repeated on 177 proteins whose structures are experimental coordinates (X-ray or cryo-EM) from the RCSB rather than AlphaFold predictions, with SS8 and SASA recomputed from those coordinates by the same pipeline. (a) Condition separation (silhouette) across depth tracks the AlphaFold result and reaches its minimum at layer 35 in both. (b) The integration index (mean pairwise CKA) increases with depth in the same way. The function channel stays orthogonal in both sets, with function-pair CKA at or below 0.12 versus about 0.99 for fused pairs. Fusion is therefore not an artifact of the predicted coordinates.

